# Differential proteome diversification in yeast populations: modes of short-term adaptation and fitness outcomes

**DOI:** 10.1101/2023.11.28.568995

**Authors:** Maor Knafo, Shahar Rezenmen, Reinat Nevo, Igal Tsigalnitski, Ziv Reich, Ruti Kapon

## Abstract

Short-term proteomic adaptations serve as an initial line of defence, allowing populations to cope with environmental changes before long-term genetic alterations occur. Using a representative set of genes, we examined how stress affects gene expression variability for different types and levels of abiotic stresses and how this influences population-level adaptation. Our data reveal that, depending on the nature of the stress, two distinct modes of response can be employed. In one, the levels of most proteins vary between individuals, leading to varied fitness levels in the population. In the other, a more limited range of expression is seen, and fitness is more even. This suggests different levels of complexity and plasticity in adaptation to different types of stress.

## Introduction

Adaptation is a fundamental process that, together with selection, shapes the evolution of a population. It is required whenever there is a mismatch between the state of a cell and the conditions in which it finds itself, a situation often referred to as stress^1–3^. Adaptation processes occur on different time scales, from the long, which allows for genetic changes, to the very short, which depends on bet-hedging and manipulation of protein levels^4,5^. Here, we concentrate on short-term adaptations that are the first line of defence in severe or unpredictable stressful situations.

As part of the tools they have at their disposal, cells can adjust the level of some of their proteins^6,7^. It is generally accepted that cells invest resources in controlling the concentration of proteins to be appropriate for their current circumstances^8–10^. However, the production of any protein involves several steps, each consisting of some stochastic elements^11,12^, that cause its abundance to vary from cell to cell^13^. This is true even within an isogenic population grown under homogeneous conditions^14^. In agreement with this notion of variation as a product of stochastic processes, the level of noise in protein amounts, irrespective of conditions or specific control circuits, has been shown to be highest for genes that are expressed at low levels and to decrease with the level of expression^6,14,15^. Indeed, previous work has shown that the level of noise in gene expression, as quantified by the coefficient of variation (CV^2^), scales as 1/p, where p is the expression level^6^.

Deviations from optimal expression can result in loss of fitness^16^ and disease^17^. On the other hand, variation in gene expression has been suggested as a bet-hedging strategy^18^, allowing cells to survive changes in conditions that entail a different set of expression levels. Thus, control of expression level is necessary, but so is variation.

The above statement stands true also under stress. However, in these suboptimal conditions, the resources available to maintain control and the returns on variability may be different. Indeed it has been shown that, when faced with extreme or unfamiliar stresses, a loss of control can ensue, which increases phenotypic variability^19,20^. This, in turn, could result in increased death rates but also has the potential to enable the exploration of various potential solutions or alternative paths to the same solution.

We wanted to address the question of how variability changes with applied stress, within the broader context of studying how adaptive mechanisms, that are rooted in single-cell behaviour, can rescue a population far from its familiar habitat. In niches inhabited by yeast, population variability has been hypothesized to be linked to survival^21,22^, so heterogeneity in gene expression in yeast is an excellent system for this investigation.

Many factors can induce cellular stress. Some are environmental and can vary significantly in type and magnitude^23,24^. Others are inflicted by biological entities in the struggle for natural resources. Existing experimental frameworks usually treat stress as a binary phenomenon, such that perturbation is either present or absent. However, there is no *a priori* reason to believe it is dichotomous and cannot vary in intensity. In this work, we treated stress as a quantifiable parameter and applied it along the full range that cells can tolerate. This required several things from the experimental system; The first was to use different types of stressful conditions for generality. Secondly, it was necessary to vary stress in increments along the entire range in which cells were viable. Finally, we needed a method to quantify the insult in a way that would allow us to compare cells undergoing different stresses.

Surprisingly, we found two different modes of adaptation, depending on the stress. Under some stresses, heterogeneity in gene expression is unaffected, hinting at similar responses across the population. On the other hand, other stressors elicit an increase in variability, which we believe reflects the availability of multiple responses that allow different solutions to the stress at hand to be formed.

## Results and Discussion

### Gene Selection and Library Representation

We constructed a representative library of fluorescently tagged yeast mutants that captures the essential features of noise in gene expression in this organism, as measured previously by others^6,25,26^, and could thus serve as a proxy for the entire proteome; We selected 52 genes that reconstitute the observed relationship between CV^2^ and average expression from a larger dataset of 2900 genes described by Newman et al (Fig. 1A). To this, we added 10 genes that are outliers to this curve^15^ as measured by Keren L et al. and five genes with well-established stress-related functions (detailed list in Supplementary Table 1).

**Figure 1.**
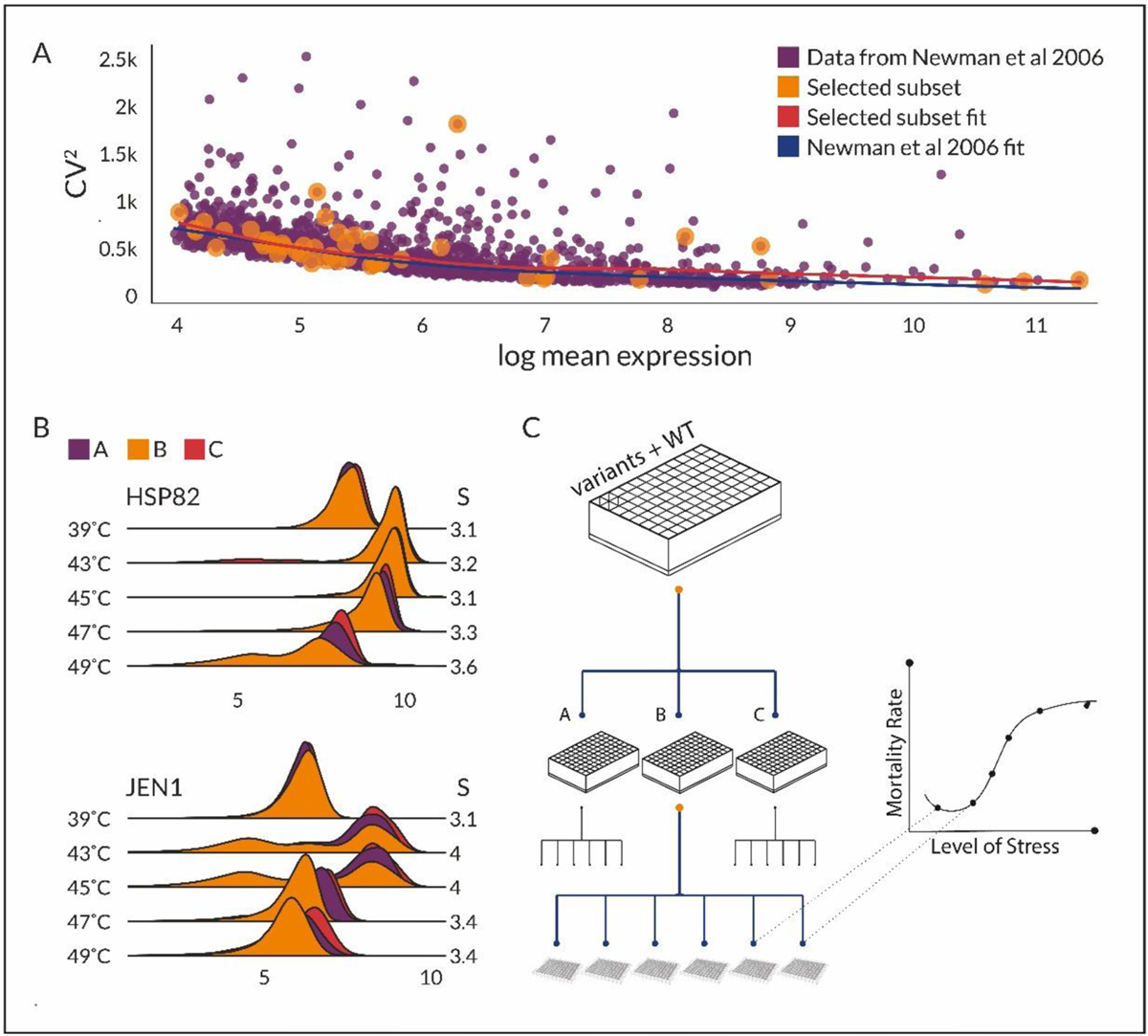
Experiment on the relationship between stress and gene-expression variability. A – CV^2^ vs log_10_(mean expression) for the original data from Newman et al. 2006 (purple), and the 52 genes we selected to represent it (orange). The blue line is the regression trend of the original data, and the red line is the regression trend for the selected genes. B - Comparison of the distribution of expression of two genes. Shown are ridge plots of the distributions of HSP82 (Top) and JEN1 (Bottom), each measured in three replicates (A, purple, B orange, and C red) at five different temperatures; right-hand values indicate the mean Shannon entropy of the distribution in bits. HSP82 presents a unimodal distribution that shifts with temperature while remaining relatively narrow up to 49°C, whereas JEN1’s expression becomes multi-modal at some temperatures. C – Experimental setup. Cells are grown overnight in three 96-well master plates, where each well in a given plate contains a single variant. Master plates are then separated into at least 6-microtiter plates, each with different stressful media/ environmental conditions, and cells are grown in them for 4 hours.

Utilizing this selection of genes, we constructed a library of 67 strains, each having one gene tagged by ymUkG1, a green fluorescent protein. These strains co-express ymUkG1 with the respective target protein, separated by a T2A^27^ ribosomal skip, so that the endogenous protein and florescent protein are expressed together, and fluorescence is proportional to protein expression level, without interfering with the endogenous protein’s function. To preserve native control and genomic environment^28^, CRISPR-Cas9^29^ was employed to place the genes at the appropriate genomic loci.

### Stress Conditions and Response Monitoring

We employed four distinct abiotic stress effectors at a series of intensities, ranging from minimal to lethal: NaCl (0.1M - 3M), H_2_O_2_ (0.1 mM-35 mM), temperature (30°C-49°C) and pH in acidic and alkaline ranges (pH 2 - pH 9) (detailed concentrations in Methods).

To avoid measuring transient responses yet prevent genetic mutations, we standardized an exposure time frame of 4 hours for each stress condition. Each strain was exposed to the full spectrum of stress conditions (31 in all, Fig 1C) and was then fixed. Single-cell fluorescence was subsequently measured in triplicate using FACS (Fig. 2, bottom). Cells with compromised outer membranes, identified using far-red Live/Dead staining (L34972 Thermo Fisher), were excluded from the analysis.

**Figure 2.**
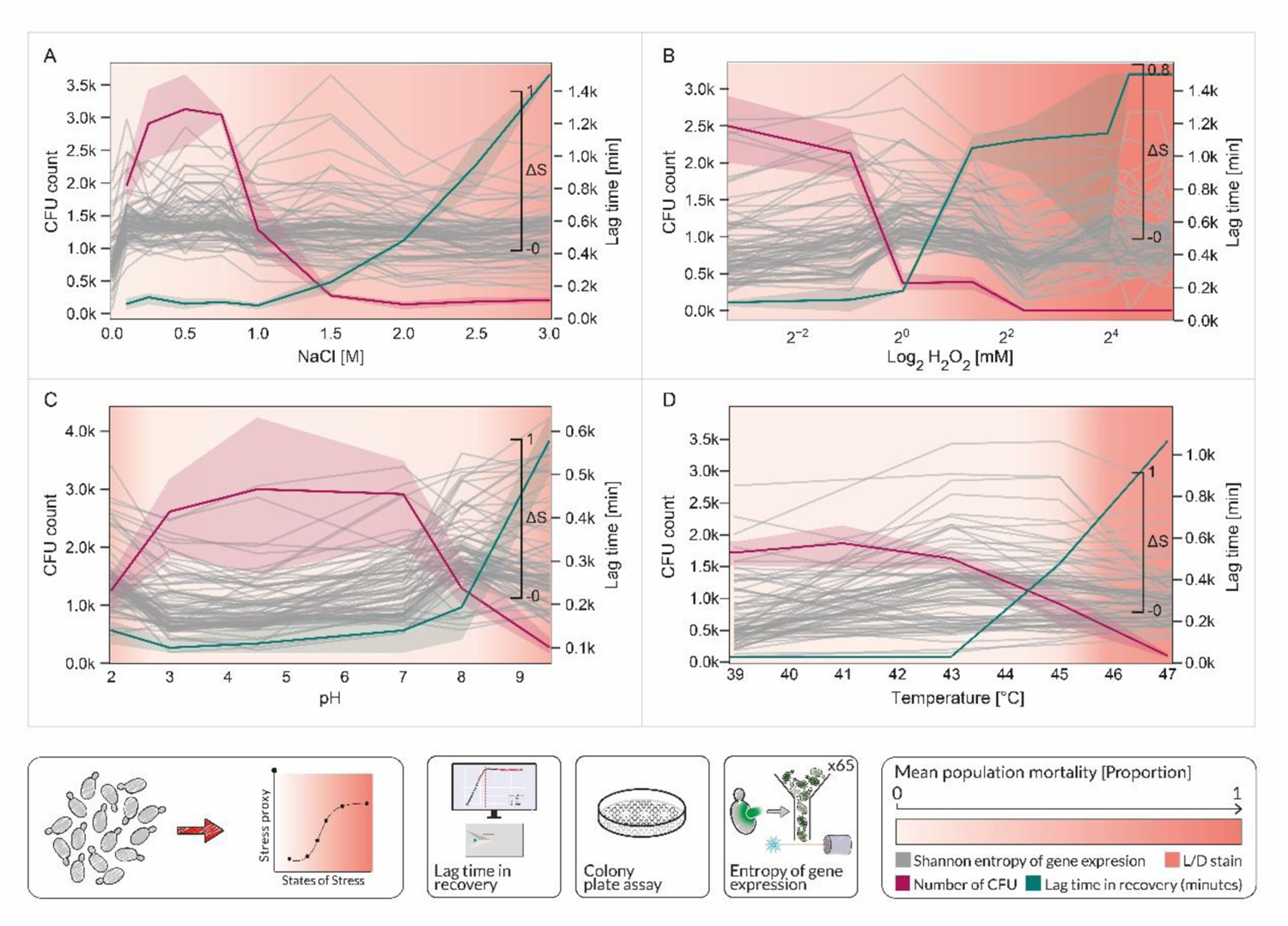
Gene expression variability as a function of stress. The upper four panels show stress and variability as a function of the various conditions applied to induce stress A - NaCl B - H_2_O_2_ C – pH, D - Temperature. Stress was quantified by the following parameters: CFU on plates (pink), lag-time in recovery (green), membrane integrity (as the intensity of the salmon background), and the variability in the expression of each gene in terms of the Shannon entropy change (grey). Data are represented as mean ± SEM. (See also Figs S6-S9 for more details). Bottom: experimental setup from left to right: WT cells are exposed to multiple levels of stress, and insult parameters (lag time in recovery, CFU number) are measured. Fluorescent reporter strains are also exposed to stress and gene expression variability (entropy of single-cell fluorescence) and the percentage of the population that is alive are measured by FACS.

Our quantification of stress is based on the state of cells following exposure, which can range from viable through growth arrest to death. We, therefore, measured three parameters: the count of colony-forming units (CFUs) that quantifies the percentage of viable cells, the lag time for recovery post-stress in conducive media, which is again indicative of growth arrest percentages but is also sensitive to how far cells are from an optimal growth state, and live/dead staining for assessing cellular membrane integrity. The latter was employed on our fluorescent strains, while the former two were quantified in populations of wild-type cells. Together, these metrics offer a more complete view of the impact of stress, a phenomenon inherently challenging to quantify. Fig. 2 delineates the stress intensity gauged using our three chosen metrics across all tested conditions, effectively mapping the spectrum from minimal impact to death.

### Entropy as a Measure of Gene Expression Variability

Gene expression variability can be characterized in various ways, depending on the parameters one wishes to quantify and the nature of the data distribution. CV^2^ effectively quantifies the breadth of the data and its distribution around the average for parametric distributions with a single peak. However, 86.610%±0.001 of the distributions we obtained were multi-peaked (Supplementary S2), which makes the CV^2^ strongly influenced by the distance between peaks rather than their individual widths. Furthermore, evaluating the expression patterns of our 67 genes revealed that 99.60%±0.03 of the distributions were non-normal (D’Agostino test at a stringent α<0.001, see Methods, Supp Table 2), and that the residuals from a normal distribution deviated significantly for all of the replicates (Supp S3).

Since our main interest is the degree to which a population explores gene expression levels, variability should chiefly quantify the partition of expression levels between possible values. A natural parameter for measuring this is the Shannon entropy, which quantifies the range and evenness of a distribution of values. The more values of expression are possible, and the more equally likely they are to occur in a population, the less well-defined the population expression level is and the higher the entropy. The calculation consisted of binning the fluorescence data, normalizing by the overall number of cells to produce a distribution of probabilities of a particular fluorescence value *p*(*f*), and calculating the Shannon entropy, *H*[*p*(*f*)] = ∑*_fmin_*^*f*max^ *p*(*f*) · log[*p*(*f*)], where the limits of the sum *f*_min_, *f*_max_, the number of bins (120) and the overall number of cells (10,000) are the same for all conditions.

Fig. 2 shows the entropy change of gene expression for each gene we measured (grey lines) as a function of the type and intensity of stress experienced by the population. As an example, consider JEN1 (distributions shown in Fig 1 B). The entropy of expression in basal conditions is 3.1. When stress is induced in the form of heat shock at 43°C, expression becomes double-peaked, and entropy rises to 4 (with a change of 0.9 bit). When the double peak distribution collapses again to a slightly wider single peak (47°C), the entropy drops to a value of 3.4 (with a change of 0.6 bit). At the same stress level, HSP82 is activated, as can be seen by the rise in the mean expression, but the distribution remains tight and maintains the same population variably at an entropy of 3.2 (0 bit).

### The relation between gene expression variability and insult levels depends on stress type

Increasing levels of NaCl From 0M to 1.5M (Fig. 2A and Supp S6) induce a drop of CFU from 3,000 colonies to several dozen, accompanied by a sharply rising lag time that reaches more than 1400 minutes, together indicating intense stress. At the same time, the entropy of gene expression rises at the very lowest level (0.1 mM) but does not change much for most genes after that. So, while all values of entropy are significantly different from basal conditions (Kruskal-Wallis with Dunn’s post hoc test p<0.0001), they remain within a narrow range of ± 0.25 bit of entropy compared to basal conditions. Similarly, exposure to hydrogen peroxide (Fig. 2B and Supp S7) leads to a drop in CFU from 2500 to 500 in the range of 0-10mM H_2_O_2_, mirrored by a significant increase in lag time in recovery from 90 minutes to above 1000 minutes. This was accompanied by an entropy rise that varied between genes but was within the range 0.3-0.4 bits compared to basal conditions. Entropy for exposure to hydrogen peroxide in a concentration above 10mM drops again when the population is no longer viable, as indicated by the fact that no colonies form after this kind of exposure.

A markedly different picture is seen when cells are challenged with increasing temperatures (Fig. 2C and Supp S8). Here, the correlation between the different stress parameters is lost. First, at 43°C, the number of colonies formed starts to drop, while the lag time in recovery remains unchanged, and the cells are intact. This could indicate that while almost a third of the cells cannot divide, there remains a population that continues to divide at an unchanged rate, even at elevated temperatures. These findings coincide with known responses to heat shock in which chaperons sustain cells despite severe stress, while allowing variable phenotypes to arise. Indeed, concomitant with this rise in temperature, a rise in gene-expression entropy is observed for a significant number of measured proteins, with the mean increasing by 0.5 bits, indicating increased variability. The almost global rise in entropy indicates that most proteins measured are distributed more widely under temperature stress, suggesting the prevalence of multiple, perhaps adaptive, phenotypes in the population when stress increases. Between 43°C and 45°C, the lag time in recovery rises to 500 minutes, the number of colonies drops to below 1000 and we begin to see membrane perforation, together indicating that more and more cells have been eliminated by selection. At the same time, the entropy of gene expression for most genes remains the same, either because the remaining cells, albeit being a smaller population, continue to explore even further or because selection acts evenly on the states reached before. Above 45°C, CFU drops significantly, the membranes of more and more cells are compromised, and the lag time in recovery is extreme and cannot be considered as part of the first line of adaptation. At this range, the entropy of gene expression declines. All these point to stricter selection.

A similar type of complex behaviour, and to an even larger degree, is observed when the population is challenged with excessively alkaline or acidic conditions (Fig. 2D and Supp S9). Between pH 3 and pH 7, all parameters indicate no or little stress and the entropy of gene expression for almost all genes is unchanged. Between 3 – 2 pH and 7 – 8 pH, lag time rises to around 200 minutes, CFU drops below 1000, membranes mostly remain intact, and the entropy of gene expression rises by 0.5-0.75 bits when compared to basal conditions for almost all genes. In these measurements, gene expression entropy is thus coupled to pH, giving the curves a concave shape, with significant differences from basal conditions (pH 4.5) for both high and low levels. At pH 9 a considerable drop in colony numbers (from 4,000 at pH 4.5 to several CFU in harsh conditions), an increase of lag times to above 500 minutes and membrane perforation of 70% of the population is observed, concomitant with a decrease in gene expression entropy.

Looking at all of our results together, we see that temperature and pH stress responses are different than those of NaCl and, to a lesser extent, of those of H_2_O_2_. Subjecting cells to low to moderate temperatures and pH levels results in a smaller number of viable cells that, however, remain intact and can resume growth when transferred to non-stressful conditions (Fig 2. C, D). At the same time, gene expression variability increases with the entropy rising by 0.6 bits on average for pH and 0.5 bits on average for temperature stress. In contrast, the response to NaCl stress includes a concomitant change in all stress measurement parameters but a small overall response in terms of gene expression variability, which rises by 0.07 bits for NaCl (Fig. 2A) and 0.25 for H_2_O_2_ (Fig. 2B).

We note that the behaviour of genes that were chosen because they display unusually high variability under conducive conditions is the same as those that lie on the general CV^2^ vs average expression curve, as is that of stress-related genes (See Supp S6-S9).

The disparity between conditions in the association between stress and gene expression variability led us to hypothesize that there are two general modes of stress response. One in which individuals differ in the expression levels of many of their genes, generating a large array of different states in the population that yield different fitness, and another, in which few genes react to the stress and most surviving cells are in the same state. In cases where the first mode is utilized, the surviving culture will comprise cells originating from the fittest of the many tried adaptations. The second will consist of a random sample of cells that share approximately the same fitness.

To test our hypothesis, we monitored the dynamics of adaptation of populations under extreme representatives of the two modes we observed, pH 8 and 1M NaCl using gUMI-BEAR, a DNA barcoding system we have developed^30^.

### Lineage-tracking experiments show that individual cells’ NaCl stress response is generic whereas pH stress induces different adaptations

Sequencing barcoded populations provides the frequency of lineages in a culture, and time-dependent changes in the frequency of specific lineages can be used to infer their fitness: lineages that are fit relative to the population will grow in frequency, whereas those that are less fit will decrease in numbers. Thus, the distribution of lineage frequencies in a population harbouring fitness differences will become skewed and the number of lineages will decrease. Populations with relatively equal fitness will keep most of their lineages until processes such as neutral drift start becoming significant.

We exposed repeats of a barcoded population to NaCl (1M, three repeats) and pH stress (pH 9, and pH 2, each in three repeats) and for each experiment, we included a control (YPD, three repeats) in naïve media. We followed the clonal makeup of these populations during 24-hour by retrieving samples every hour and later sequencing them.

When exposed to NaCl (1M), we observe that the number of lineages, as well as the distribution of relative lineage sizes (Fig. 3A Red & Green), remains unchanged with time. The latter is also seen quantitatively in Fig. 3C where the change in entropy of the distribution of lineage sizes is plotted as a function of time. The fact that the distribution of lineages does not become narrower or significantly skewed after 24 hours shows that the division rates of all lineages are similar. At the same time, our previous CFU quantification showed that 50% of the cells die at this concentration (see CFU in Fig. 2A). Together, these two observations indicate that all lineages lose cells, and they do so at a similar rate. These results, and the fact that we did not observe, other than the initial rise with minimal exposure to salt, a rise in entropy in gene expression, suggest an even proteomic response shared by all lineages, which have the same fitness and likelihood of survival, randomly distributed across the population.

**Figure 3.**
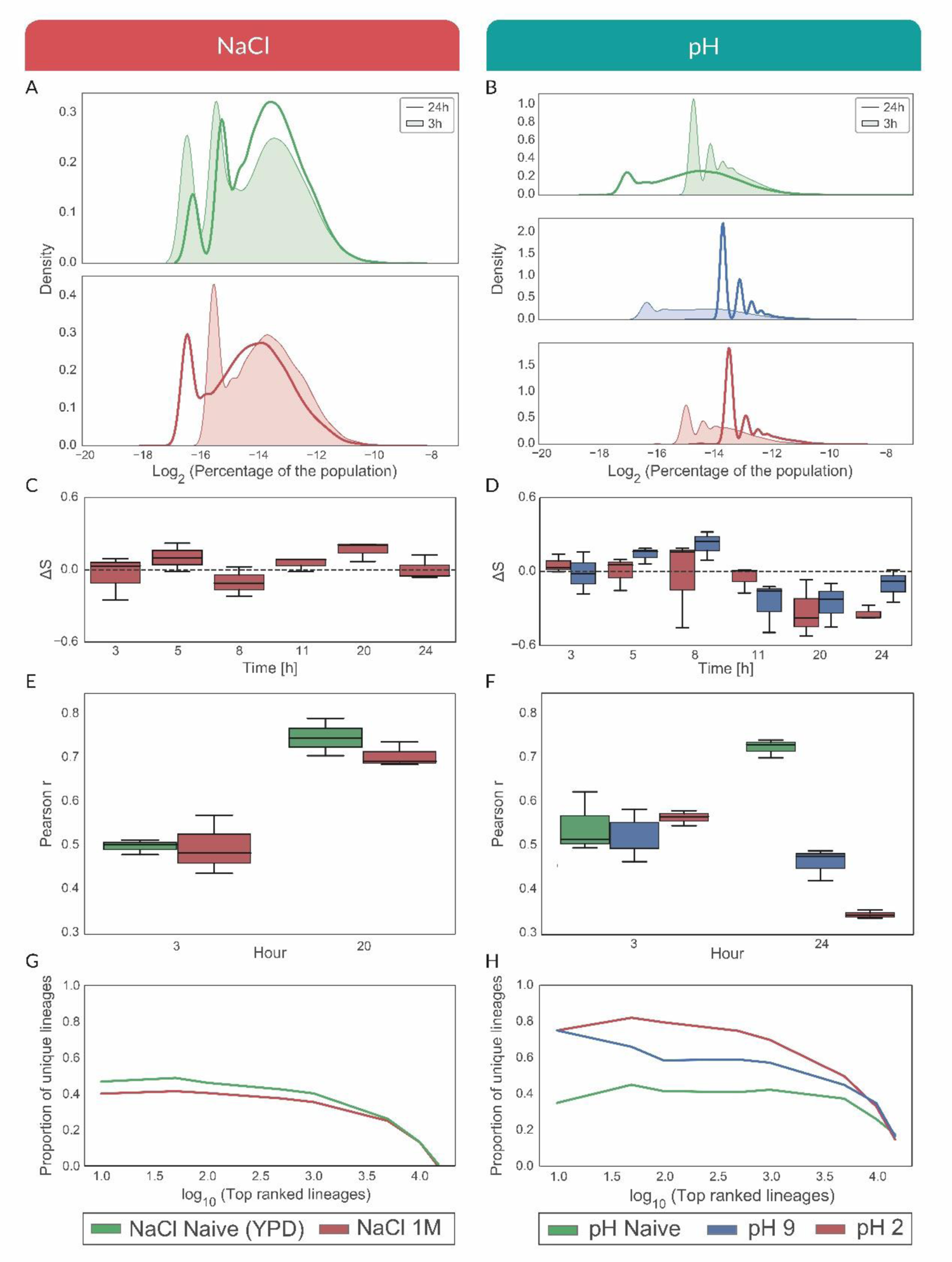
Lineage tracking experiments show differences in the modes of adaptation to NaCl (left), and pH (right). In all graphs, green is an unchallenged population, and red (NaCl on the left and pH 2 on the right) and blue (pH 9) indicate a challenge. A, B - The distribution of lineage sizes for unchallenged cells (green), NaCl challenged cells (A, red), low (B, red) and high (B, blue) pH challenged cells at the beginning (thin line, shaded) and end (heavy line) of the experiments. C, D - Change in entropy of lineage distributions (ΔS) between challenged replicates as a function of time. E, F Pearson correlation between biological replicates at the beginning and end of the experiment. G, H - Proportion of lineages that are unique to a repeat for increasing numbers of top-ranked lineages.

Our gUMI-BEAR methodology allows us to produce identically barcoded libraries^30^. When comparing the lineage composition of populations between repeats, we found that it is highly similar for both NaCl treatment and for the unchallenged populations (Fig. 3E). Over half of the top three thousand most successful lineages are shared by all repeats (Fig. 3G, Red, Fig. 3G, Green). We therefore believe that either all cells experiencing NaCl stress adapt in a very similar manner or that no adaptation occurred during the experiment, leaving the structure of the population relatively constant.

The response to pH 2 and pH 9 stress is entirely different. Around 8 hours into the stress, the number of lineages decreases, and the distribution of lineage frequencies becomes tighter with the representation of the highest-ranking lineages increasing (Fig. 3B). This would indicate the development of fitness differences that are heritable within lineages. At the same time, the composition of the population (Fig. 3D Red & Blue) for both stressful pH conditions and the identity of successful lineages (Fig. 3F) differs between repeats, as indicated by the decreasing correlation and an increasing number of unique successful lineages between replicates (Fig. 3H). Therefore, the relative fitness gained by specific lineages is the result of a stress-induced adaptation, occurring during the experiment and not of processes that may have occurred before the library was divided between repeats. Combining this with the rise in gene-expression entropy, that we observed previously, suggests that multiple pathways are available to each cell in the population to combat pH stress and that these are “turned on” by stress and lead to the ability to survive, albeit at different fitness levels.

## Discussion

In our experiments, we observed two distinct modes of adaptation to stress in yeast populations (Fig. 4A). In the first, variability in gene expression between individual cells remains unchanged for most of the genes we tested, and clones exhibit little fitness differences. Thus, selection acts equally on *all* lineages in a population, and the overall structure of the population does not change under stress. This type of response was brought on by exposure to NaCl and, to a lesser degree, to H_2_O_2_. On the other hand, pH and temperature stress led to a significant rise in the variability of expression of almost all genes tested and to distinct lineages gradually gaining fitness over others. This suggests that adaptation to these stresses manifests as individual cells adopting one of many different states of the proteome, each constituting a different combination of gene-expression patterns that lead to varied fitness values. In this mode, the fitter clones become more dominant with time. The fact that the clonal identity of the more successful lineages varied between repeats of the experiment points to adaptation being induced by the stress and not resulting from pre-existing differences between lineages. Thus, the possibility for all solutions must have existed in each clone.

**Figure 4.**
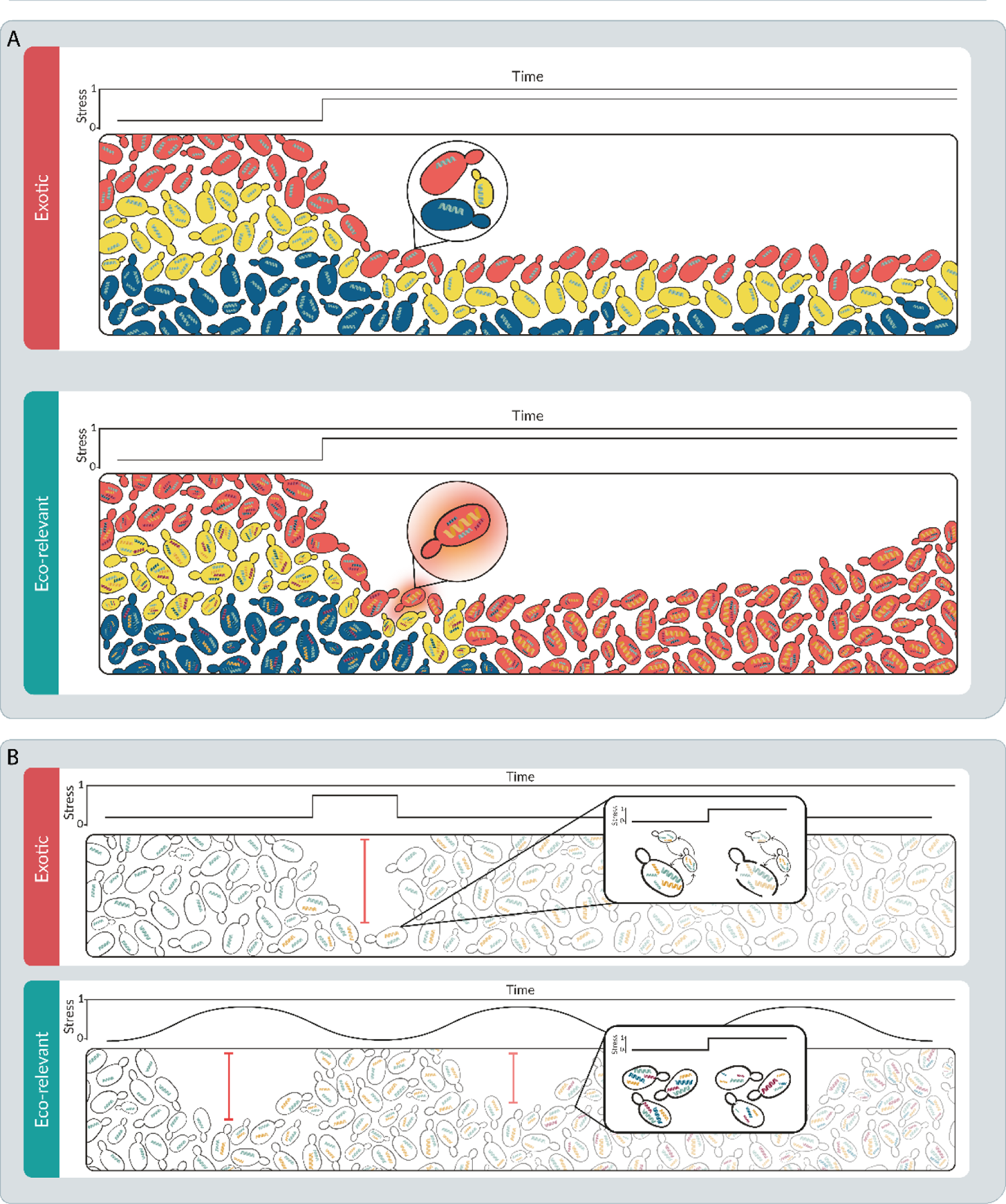
Richness in Stress Response Strategies. A- (Top) When exposed to NaCl stress, the population exhibits a uniform response, with different lineages, all possessing a single expression level (inset: uniform single green protein across lineages). As a result, the diversity of both gene expression and lineages remains constant despite strong selection. (Bottom) In contrast, pH stress leads to exploration in gene-expression values, increasing its variability, ultimately leading to adaptive events in specific lineages, such as the red lineage (inset: red cell enriched with yellow protein), causing it to outgrow others, thereby reducing lineage diversity. B Hypothetical trajectory that might lead to generic vs robust proteomic response (Top) Infrequent historical exposure to exotic stressors results in a lack of specialized responses. Cells navigate through a phenotypic landscape, with survival determined randomly (inset: random survival in a phenotypic landscape). (Bottom) Frequent historical exposure to stressors equips cells with multiple adaptive solutions (inset: diverse cellular responses to stress), leading to a resilient and robust network of coping mechanisms.

How can we explain such differences in stress-adaptation mechanisms that ultimately lead to similar degrees of survival? One possibility is that the type of mechanism available to cells depends on the degree of prevalence of the stress. Stresses encountered by yeast range from infrequent stresses, such as NaCl, seldom encountered by yeast populations in Nature and highly toxic to them^31–33^, to ecologically relevant stresses that often perturb yeast populations, such as pH or temperature fluctuations. It is possible that the first mode of adaptation we observed, in which all cells take on the same solutions and obtain similar fitness, is employed when an unfamiliar stress is encountered^34,35^. On the other hand, when assaulted with familiar stress, the cell has a comprehensive arsenal of proteins^36^ whose expression can be modified at its disposal, resulting in multiple, distinct, single-cell events, each with its own fitness^37^.

Both modes of adaptation can account for stress resilience, but we believe that the increased variability resulting from the second course can serve as a form of biological plasticity, enabling the rapid reconfiguration of phenotypes to withstand environmental insults.

Based on our results, we speculate that the aforementioned plasticity can be formed by repeated cycles of selection, expansion under stress and relaxation under optimal (or different) conditions, where each cycle can lead to integration of new stress-alleviating mechanisms (Fig. 4B). This suggestion is in line with recent theoretical work by Gómez-Schiavon and Buchler^38^, which shows that under repeated and frequent environmental change, populations tend to adopt a solution based on transitioning a constitutive gene to an epigenetic switch. This type of solution is preferred because it reduces the time necessary to adapt to changing conditions (as compared to adaptation by mutation) while maintaining robustness in the face of biochemical noise. The fact that some of the gene expression patterns we found are multi-modal lends further support for this proposal.

Another support is provided by a study by Yang and Tavazoie^39^, where they “familiarize” yeast to higher-than-usual, lethal concentrations of ethanol by implementing a frequent cycle of exposure-relaxation, similar to that described above. The adapted strains integrate many additional genes into the global response to severe ethanol exposure compared to the native strain. The upregulated and downregulated proteins are diverse in function and responsibilities, from specific physiological to general stress responses.

It is interesting to note that the most differentially expressed genes in the Yang study were of unknown function and, as such, raise the intriguing notion that previously unfamiliar stress can eventually be alleviated by harnessing functions of various origins, adding these to the cell’s network of stress response. The more frequent the stress, the more cell machinery it harnesses to accommodate it, yielding many pathways for adaptation in unique, novel, and creative ways. This hypothesis could be tested experimentally by repeating unfamiliar stresses at varying intervals and observing the appropriation of genes towards novel solutions.

## Acknowledgments

The authors wish to thank Tzachi Pilpel, Roni Paz, Orna Dahan, Jonathan Nutkiewitz, and Nathalie Balaban for helpful scientific discussions and Shira Holand for the scientific graphics.

## Author contributions

Conceptualization M.K., Z.R. and R.K. Methodology M.K., R.N., Z.R and R.K., Software, M.K., Validation, M.K., Formal Analysis, M.K., S.R., I.T., Investigation, M.K., S.R., I.T., Resources, M.K. Data Curation, M.K., Writing, M.K., R.N., Z.R. and R.K., Visualisation, M.K. and R.N., Supervision, R.K.

## Lead contact

Further information and requests for resources should be directed to and will be fulfilled by the lead contact, Ruti Kapon

## Methods

### Materials availability

Further information and requests for resources and reagents should be directed to and will be fulfilled by the lead contact.

### Experimental model

BY4741 strain *Saccharomyces cerevisiae* (*S. cerevisiae*) haploid cells were obtained from the Weizmann Institute core facility unit and kept at −80°C in a 50% glycerol and 50% YPD (Yeast Peptone Dextrose) solution.

## Method details

### Variant construction

Variants were produced using CRISPR/Cas 9 technology. The donor DNA construct consisted of five key elements (see map in Supplementary Information S1): (1) 100 bp left homology arm (LHA) to guide the construct to the correct genomic location, aligning with the region upstream of the gene of interest. (2) *ymUkG*1 - The fluorescence reporter gene used for quantification. We chose ymUkG1 as a fluorescent tag because of its brightness and the high correlation of its expression with known promotor activity^40^. (3) *T2A* - A ribosomal skip sequence which ensures the ymUkG1 fluorophore is translated as a separate protein that does not interfere with the function of the native protein. (4) NGS-ready barcode - A unique identifier that enables the identification of variants in sequencing. (5) 100 bp right homology arm (RHA) - This sequence aligns with the region downstream of the gene of interest, further ensuring correct integration. The constructs were synthesized by Twist Bioscience and subsequently PCR-amplified (Initial denaturation: 98°C for 1 minute. 30 cycles of: Denaturation: 98°C for 30 seconds, Annealing: 60°C for 15 seconds, Extension: 72°C for 1 minute. This is followed by a final extension at 72°C for 5 minutes.) in Kapa Master Mix (Kapa Biosystems, Catalog No. KK2602) using generic primers (F-GAAGTGCCATTCCGCCTGACCT, R-CACTGAGCCTCCACCTAGCCT).

A gRNA for each gene-fluorophore pair was flanked by generic integration sites and cloned into a pCAS plasmid using restriction-free (RF) cloning (in Kapa Master Mix) according to the following protocol ^29^: Initial denaturation: 98°C for 1 minute. 30 cycles of: Denaturation: 98°C for 30 seconds, Annealing: 58°C for 1 minute, Extension: 72°C for 10 minutes. This is followed by a final extension at 72°C for 10 minutes. The plasmid is held at 4°C. The integrity of the plasmid and the correct insertion of gRNA sequences were validated by PCR, using the primer 5′-CGGAATAGGAACTTCAAAGCG-3′.

Cells (90 μl of 1 OD 10X concentrated culture) were prepared for transformation by resuspending in 900 µL of PLATE solution (PEG 2000, lithium acetate, Tris EDTA buffer) with 10 µL of single-stranded DNA from salmon sperm (Sigma-Aldrich, Catalog No. D7656-1ML). For each gene, 1µg of the corresponding pCAS plasmid and 5µg of the donor DNA were added. The cells were then incubated for 30 minutes at 30°C and subjected to a 15-minute heat shock at 42°C. Following heat shock, the cells were pelleted and resuspended in 250 µL of YPD for a three-hour recovery period at 30°C without shaking. Then, the cell suspension was diluted with 2 mL of YPD supplemented with G418 (Sigma-Aldrich, Catalog No. A1720), 200 mg/L, to select cells that received the pCAS plasmid. The mixture was incubated for 48 hours at 30°C, after which ymUkG1-positive cells were sorted using the BD FACSMelody™ Cell Sorter system (BD Biosciences).

The sorted cells were plated on YPD agar plates (1% agar) supplemented with G418 (200 mg/L) to select cells that have the pCAS plasmid. After 48 hours of incubation at 30°C, colonies were selected, and Sanger sequenced Cells were kept in a glycerol stock at −80°C.

### Measurement of variability under stress

Preparing yeast cells for stress measurements began with streaking a frozen aliquot of each variant on a YPD plate. A single colony from the plate was inoculated into 3 ml of YPD medium and grown overnight. The overnight culture was then transferred into 100 ml of fresh YPD medium and allowed to grow for 12 hours until it reached OD 1 at 600nm.

Three 96 deep-well master plates, labelled A, B, and C, were prepared by placing 150 µl of each variant in a different well. To control for the effects of well location in further experiments, the allocation of the variants in the wells was shuffled between master plates (see Table 1 in the Supplementary Information).

### Preparation and incubation of master plates before stress exposure

400 µl of YPD was added to each well in the master plates. These plates were incubated overnight in the HEIDOLPH-1000 TITRAMAX, set to 900 RPM and 30 °C. This step was repeated three times, each performed in one of the three biological replicates.

### Preparation of stress-inducing media

Media for inducing the different stress conditions were prepared and 135 µl of the resulting stressful media was placed in each of the wells of a 96-well plate as follows: For NaCl stress: NaCl (J.T Baker, Cas: 7647-14-5) was added to YPD to final concentrations of 0.1 M, 0.25 M, 0.5 M, 0.75 M, 1 M, 1.5 M, 2 M, 2.5 M, and 3 M. For pH stress: YPD was buffered using K_2_HPO_4_ (J.T Baker, Cas: 7758-11-4) to a final concentration of 150 mM. The pH was adjusted to pH 2, pH 3, pH 4.5, pH 7, pH 8, and pH 9 using NaOH or HCl. For Hydrogen peroxide (H_2_O_2_) stress: Hydrogen peroxide 30% (Bio-Lab, cat 000855032300) was added to YPD to achieve concentrations of 0.1 mM, 0.5 mM, 1 mM, 2.5 mM, 5 mM, 15 mM, 20 mM, 30 mM, and 35 mM. For temperature stress: A range of temperatures, which included 30, 39, 41, 43, 45, and 47 °C were set in a BINDER 9020-0313 incubator.

### Application of stress conditions

The overnight cultures from the three repeats were diluted 10X and 15 µl of each one was distributed into one of the plates containing either a stress-inducing media (NaCl, pH, and H_2_O_2_ stresses) or standard media (for temperature stress). The stressful incubation procedure was performed three times, once for each of the three biological replicates.

### Incubation under stress conditions

The diluted cultures were incubated for 4 hours under the same conditions as the overnight incubation in the HEIDOLPH-1000 TITRAMAX, set to 900 RPM at 30 °C (or the predetermined temperature levels for the temperature stress condition).

### Live/Dead staining and fixation

Cells from each of the 96-well microtiter plates were resuspended in 500 μl of the staining solution (Thermo Fisher cat - l34972) diluted 1:1000 in PBS), incubated for 15 minutes at room temperature, centrifuged at 1500g for 2 minutes, and the supernatant was discarded. The cells were then washed with 150 µl PBS. Cells were then fixed by diluting PFA FX0415 to 4% in PBS and adding the fixative solution to the palette of cells. Cells were centrifuged at 1500g for 2 minutes, the supernatant was discarded, and the cells were washed with 150 µl KPO_4_/sorbitol. This last step of centrifugation and wash was performed twice.

Fluorescence was measured in plates using flow cytometry (Attune NXT FACS machine) with excitation at 488nm and the 530/30 filter. Analysis was performed using FlowJo software (BD, Catalog No. 361071) with gates for singlets and live/dead staining (detailed FlowJo protocol in supplements). Data from live cells were extracted to CSV files, with subsequent statistical analysis performed using in-house code written in Python.

### Application and quantification of stress

#### Application of Stress

The application of stress was carried out on WT (BY4741) cells as follows: Cells were grown overnight from a single colony in 100 ml YPD media at 30°C at 220 RPM until the culture reached 1 OD. The culture was then diluted 10X to a 96-well plate containing the full range of stressful media in different wells in three biological replicates. The cells in the stressful media were then incubated for 4-hour stress exposure in the same conditions as above.

#### CFU count

15 µl of each culture was taken from the deep-well plate and was diluted 10X. 100 µl of the final dilution (equivalent to 1 µL of the original culture) was spread on agar plates. After 48-hour incubation at 30 °C, colonies on these plates were counted using a custom script in Fiji ImageJ software (see Supplemtary Information for Raw data and code) based on scanned images of the plates scanned on the GE Healthcare Epson ImageScanner III. Experiments were performed in 3 biological replicates.

#### Lag time recovery and growth curve analysis

15 µl of each culture was washed and resuspended in 135 µl fresh YPD medium. These cultures were then incubated for 24 hours at 30°C with fast (538 RPM) double-orbital shaking in the Cytation plate reader (BioTek, Catalog No. Cytation3). Recovery and growth kinetics of the treated culture were monitored, and the resulting growth curves were analysed using a custom Python script to extract key parameters, such as lag time, doubling time, and time to stationary phase (see Supplementary Information for Raw data and code). Experiments were performed in 3 biological replicates.

#### Lineage tracking using gUMI-BEAR

We utilized the gUMI-BEAR (genomic Unique Molecular Identifier Barcoded Enriched Associated Regions) method on WT yeast for lineage tracking^30^. Samples were prepared by combining seven sub-libraries resulting in a library containing 26,00 lineages. The combined library was grown in YPD overnight until it reached 1 OD in 100 ml and was then diluted to 0.1 OD. 2 ml of the resulting culture were added to each well in a 96 deep-well plate. The sample was centrifuged at 1600g for 2 minutes and resuspended in 96 well plates with relevant stressful media to expose them to a series of pH and NaCl stress levels (NaCl 0 M, 0.1 M, 0.25 M, 0.5 M, 1 M, 1.5 M and, pH 2, 3, 4.5, 7, 8, 9 and Naive), with each stressful level replicated three times in each experiment. The plates were incubated at 30°C and shaken at 200 rpm in an Inheco TEC Control 96/RS incubator (Inheco, Catalog No. 8900030). Every hour, 10% of the volume of the well was removed and preserved in DNA/RNA Shield (Zymo Research, Catalog No. R1100-250) using the Tecan Freedom EVO liquid handling system (Tecan, Catalog No. 30062315). We replenished 10% of the stressful media into each sample to account for volume reduction. Altogether, the cultures grew for 24 hours.

Samples were prepared, cleaned, and processed as described^30^. The resulting final libraries were sequenced on an Illumina NovaSeq platform at the Weizmann Institute of Science’s G-INCPM unit. Data analysis was performed using the R code provided^30^ and subsequent diversity calculation in Python (see Resource Availability for Raw data and code).

### Quantification and statistical analysis of population variation

#### Calculating entropy of gene-expression

In order to ensure consistency in entropy calculations in different gene-expression datasets, we maintained three assumptions:

#### Uniform support for measurements

Setting a consistent dynamic range for FACS measurements was crucial to have consistent probability calculations. To accomplish this, we set the autofluorescence emitted by naive cells as the lower boundary for fluorescence measurement. The highest fluorescence value measured (in the entire dataset) was used as the upper bound. *Consistent sampling depth*: For each dataset, we randomly sampled 20,000 cells to avoid overestimating low-density probabilities. *Selection of bin numbers*: To make meaningful comparisons of probabilities, it was necessary to bin the data with a consistent number of bins that also had to be large enough to represent population richness adequately but not so large so the signal wouldn’t be coherent. We calculated the entropy for an increasing number of bins and found that it plateaued at 100 bins. We, therefore, used 100 bins, evenly distributed across the dynamic range for all calculations.

Following binning, we computed the entropy using the standard formula for Shannon entropy as described in the Results and Discussion section, where the probability for each bin was calculated as the number of measurements in the bin divided by the total number of measurements.

## Data and code availability

See Supplementary Information.

