## Supplementary material for "Differential proteome diversification in yeast populations: modes of short-term adaptation and fitness outcomes": Supp

### Supplemental information

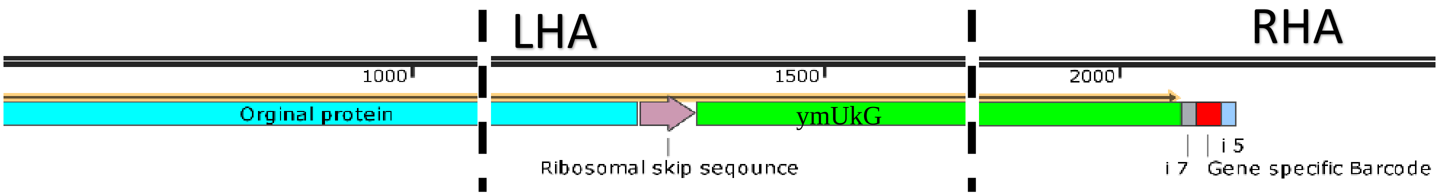


**Figure S1: Map of Donor DNA.** A donor DNA was designed for each candidate gene to be integrated into the genome at the c-terminal part of the protein using HDR (homology-dependent repair). The insert was made up of the following elements: 100bp left homology arm (LHA), T2A (ribosomal skip sequence), ymUkG1 (a GFP variant) – Gene-specific barcode- 100bp right homology arm (RHA). The sequences were ordered from Twist Bioscience and amplified using generic primers**.**

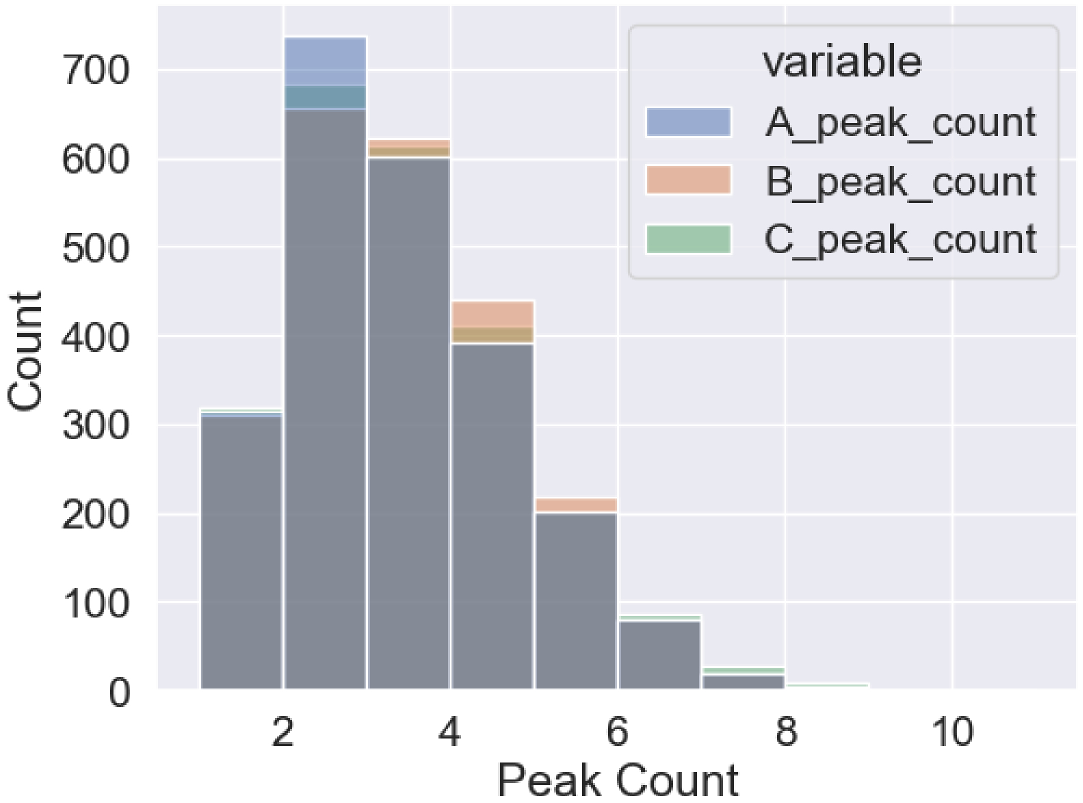


**Figure S2**: **Peak count per gene by biological replicates.** Distribution of peak counts per gene for datasets A, B, and C. The x-axis represents the number of peaks, and the y-axis denotes the count. Different colors distinguish the three repeats A- Blue, B – Orange, C- Green.

| Marked Gene | Well location in plate A | Well location in plate B | Well location in plate C |
|---|---|---|---|

| | | | |
|---|---|---|---|
| **APL2** | A1 | H12 | A10 |
| **ECI1*** | A2 | H11 | A9 |
| **GTT3** | A3 | H10 | A8 |
| **JEN1*** | A4 | H9 | A7 |
| **NAT2** | A5 | H8 | A6 |
| **PRX1** | A6 | H7 | A5 |
| **RSN1** | A7 | H6 | A4 |
| **TBS1*** | A8 | H5 | A3 |
| **YGL108C** | A9 | H4 | A2 |
| **YGR017W** | A10 | H3 | A1 |
| **ECM21** | B2 | G11 | B9 |
| **HEM15** | B3 | G10 | B8 |
| **NOP53** | B5 | G8 | B6 |
| **PYK2*** | B6 | G7 | B5 |
| **SEC5** | B7 | G6 | B4 |
| **TEF1** | B8 | G5 | B3 |
| **YJL045W*** | B9 | G4 | B2 |
| **ATP7** | C1 | F12 | C10 |
| **ENO2** | C2 | F11 | C9 |
| **HSP82#** | C3 | F10 | C8 |
| **LAT1** | C4 | F9 | C7 |
| **OSM1#** | C5 | F8 | C6 |
| **RAI1** | C6 | F7 | C5 |
| **SEG1** | C7 | F6 | C4 |
| **UBC4** | C8 | F5 | C3 |
| **NUP157** | C9 | F4 | C2 |
| **BER1** | D1 | E12 | D10 |

| | | | |
|---|---|---|---|
| **FKS3*** | D2 | E11 | D9 |
| **LNP1** | D4 | E9 | D7 |
| **PDC1** | D5 | E8 | D6 |
| **RAP1#** | D6 | E7 | D5 |
| **SFC1*** | D7 | E6 | D4 |
| **VPH1** | D8 | E5 | D3 |
| **PIR1** | D9 | E4 | D2 |
| **BUD20** | E1 | D12 | E10 |
| **GAS2*** | E2 | D11 | E9 |
| **HXK1*** | E3 | D10 | E8 |
| **MIA40** | E4 | D9 | E7 |
| **RPL27A** | E6 | D7 | E5 |
| **SLC1** | E7 | D6 | E4 |
| **YBR137W** | E8 | D5 | E3 |
| **NSA1** | E9 | D4 | E2 |
| **CBF5** | F1 | C12 | F10 |
| **HXT2*** | F3 | C10 | F8 |
| **MNN11** | F4 | C9 | F7 |
| **PEX29** | F5 | C8 | F6 |
| **RPM2** | F6 | C7 | F5 |
| **SLF1** | F7 | C6 | F4 |
| **YBR184W*** | F8 | C5 | F3 |
| **ICE2** | F9 | C4 | F2 |
| **WT1** | F10 | C3 | F1 |
| **CDC34** | G1 | B12 | G10 |
| **GPT2*** | G2 | B11 | G9 |
| **HXT7*** | G3 | B10 | G8 |

| | | | |
|---|---|---|---|
| **PGU1*** | G5 | B8 | G6 |
| **RPN5** | G6 | B7 | G5 |
| **STE20** | G7 | B6 | G4 |
| **YKL071W** | G9 | B4 | G2 |
| **WT2** | G10 | B3 | G1 |
| **CIT3*** | H1 | A12 | H10 |
| **GRE2** | H2 | A11 | H9 |
| **IES4** | H3 | A10 | H8 |
| **PIN3** | H5 | A8 | H6 |
| **STL1*** | H7 | A6 | H4 |
| **YEH2** | H8 | A5 | H3 |
| **YAL053W** | H9 | A4 | H2 |
| **WT3** | H10 | A3 | H1 |

**Table T1**: **List of genes used in the experiment and their placement in the 96-well plates in the different repeats.** Left column gives the list of 67 genes used to represent the proteome in our experiments. Noisy genes are marked in $^*$, and stress genes by $^\#$. The right three columns indicate where we placed mutants with the specific gene conjugated to the ymUkG1 in the three 96-well plates that formed the repeats in the experiments. The location of mutants within the plates was shuffled to address artifacts that could result from position.

#### *Characterizing gene expression distributions*

We wanted to characterize the distributions to help us determine how best to analyze the gene expression data. Because we are mostly interested in the shape of the distribution and its closeness to normal, we chose the D'Agostino test. We applied it to the data obtained from each replicate, using a highly conservative alpha level ($\alpha$<0.001) to minimize the risk of wrongfully rejecting the assumption that the data is normally distributed. Even under this stringent statistical threshold, our results revealed that the expression levels of almost *none* of the genes displayed a normal distribution.

| **Biological replicate** | **Proportion passed the D'Agostino test** |
|---|---|

| | |
|---|---|
| A | 0 |
| B | 0.006 |
| C | 0.005 |

**Table T2: Proportions of genes from each biological replicate (A, B, and C) that passed the D'Agostino test for normality.** Note the extremely low proportions of genes that passed the test across all replicates.

To obtain a quantitative estimate of how far the distributions were from normal, we compared the measured expression distributions to a calculated normal distribution with the same mean and standard deviation using Earth Mover's Distance (EMD). The EMD values for each gene ranged from 0.1 to 1.4. Median values for repeats A, B, & C were 0.133, 0.131, and 0.129, respectively. Compared with 100,000 pairs of Gaussian distributions with means and standard deviations within our data range, their EMD values centered around 0.014 with a width of 0.07. Figure 3D illustrates the EMD distribution for all genes and the Gaussian samples. A Kolmogorov-Smirnov test showed a p-value of <0.001 for all replicates (see Table S3).

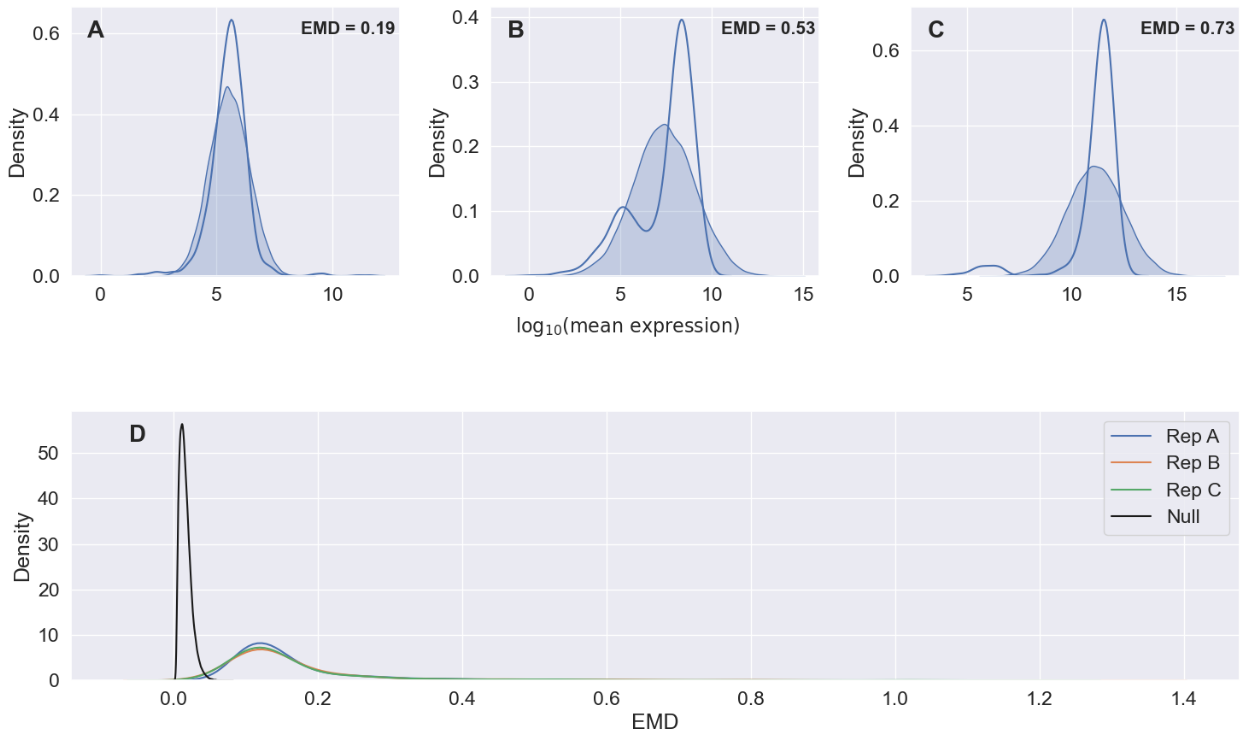


**Figure S3: Earth Mover's Distance (EMD) analysis. Examples of distributions of expression levels of 3 genes (**A) NSA1 pH4.5 replicate A, (B) PRX1 at pH 9.5 replicate B, (C) PDC1 at 43 43° C replicate B compared to a normal distribution with the same mean and STD. The calculated EMD (A) 0.19, (B) 0.53, (C) 0.73. (D) The distribution of EMD values for all genes in each biological replicate (A, B, and C) was compared against a null distribution. The Null distribution was created by performing 100,000 EMD measurements between two Gaussian random samples who's average, and width were drawn from the same Gaussian distribution.

| Replicate | KS Statistics | p-value |
|---|---|---|
| A | 0.524 | <0.0001 |
| B | 0.523 | <0.0001 |
| C | 0.523 | <0.0001 |

**Table T3: Kolmogorov-Smirnov test results comparing each biological replicate's EMD distribution to the null distribution.** The p-values are less than 0.001 for all biological replicates, indicating that the EMD distributions of the replicates are significantly different from the null distribution.

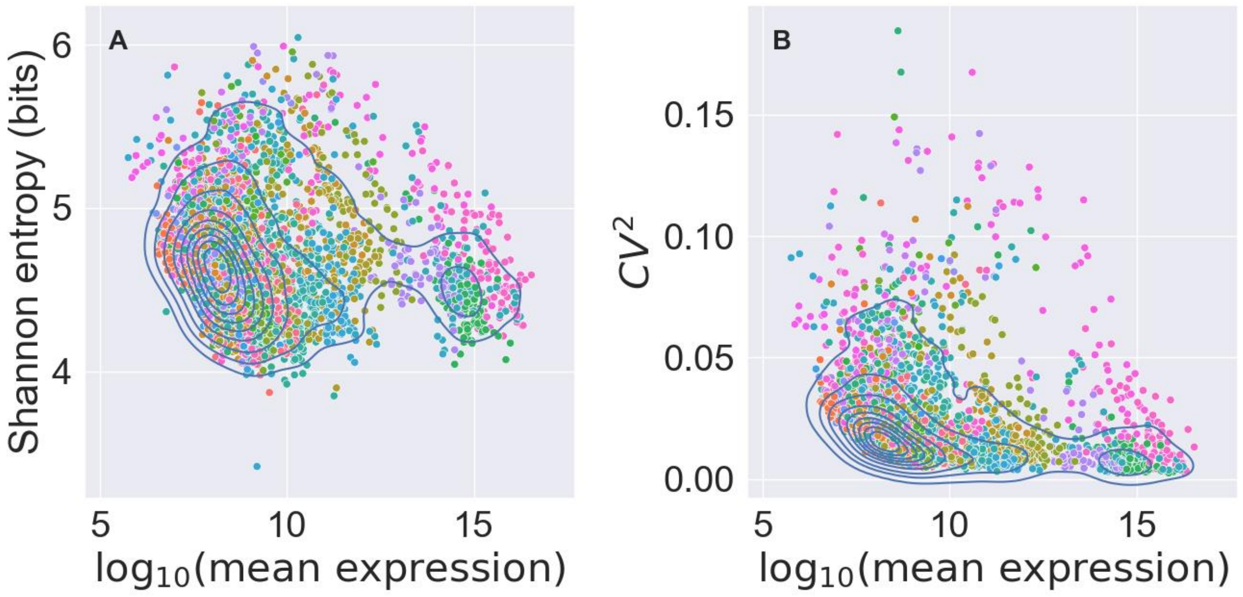


**Figure S5**: **Gene expression variability vs average expression for the entire dataset as measured by entropy and coefficient of variation. (**A) Density plot illustrating the relationship between Shannon entropy and $\log_{10}$(mean expression) for various genes. Each point represents a gene, points are color-coded based on the condition under which they were measured. (B) As in previous studies(1–3), the density plot shows CV2 goes down with $\log_{10}$(mean expression).

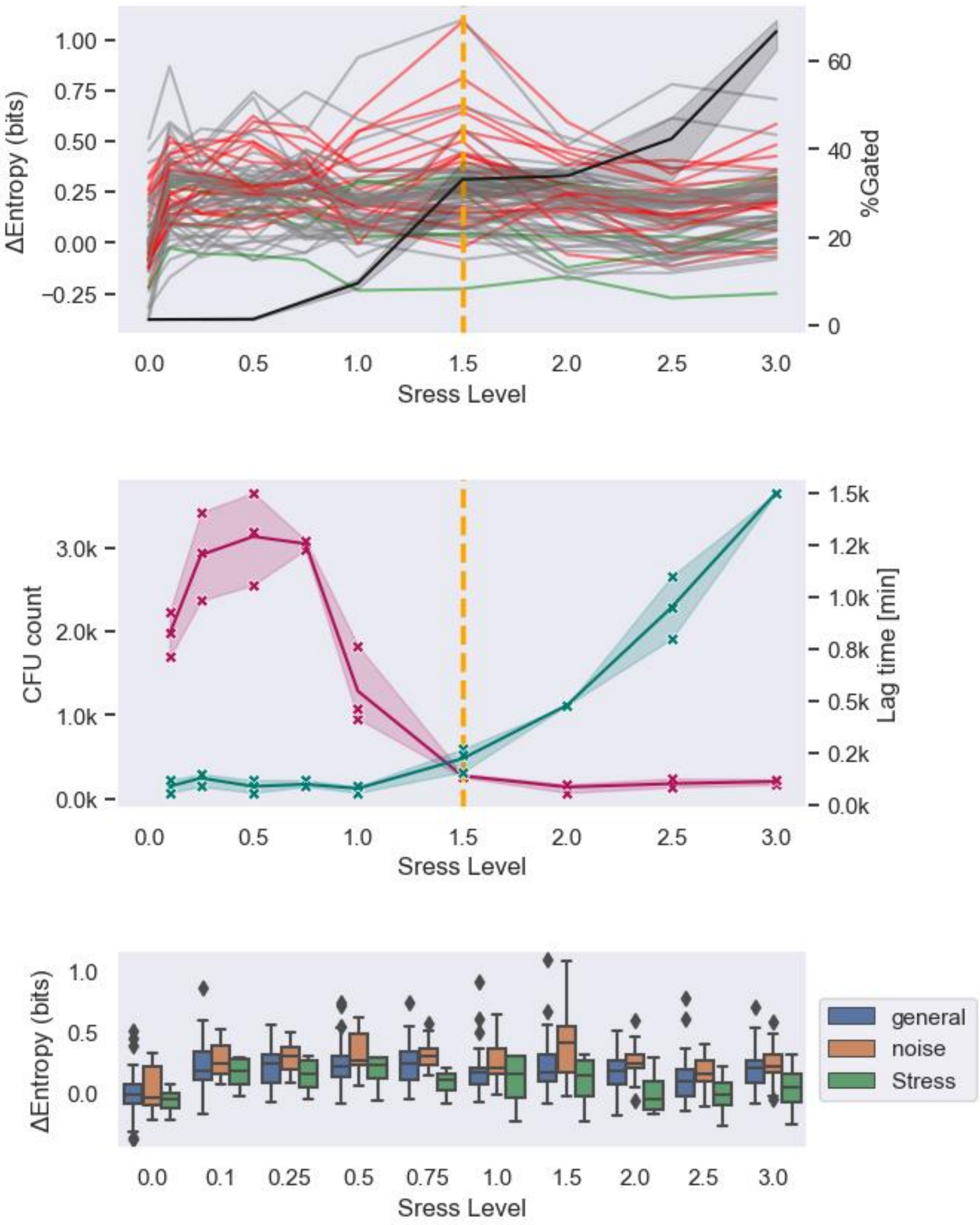


**Figure S6: Entropy of gene expression and stress parameters for different concentrations of NaCl.** The top panel shows the entropy of gene expression for all genes, where "noisy" genes are shown in red and stress genes in green. The black line shows the % of dead cells as measured using live/dead far-red stain. The middle panel shows stress quantification via CFU and recovery lag time. The dashed orange line signifies the end of the physiological range according to Yeast Stress responses (4). The bottom panel shows the entropy distribution data as the box plots. Notice that all gene groups display a similar trend.

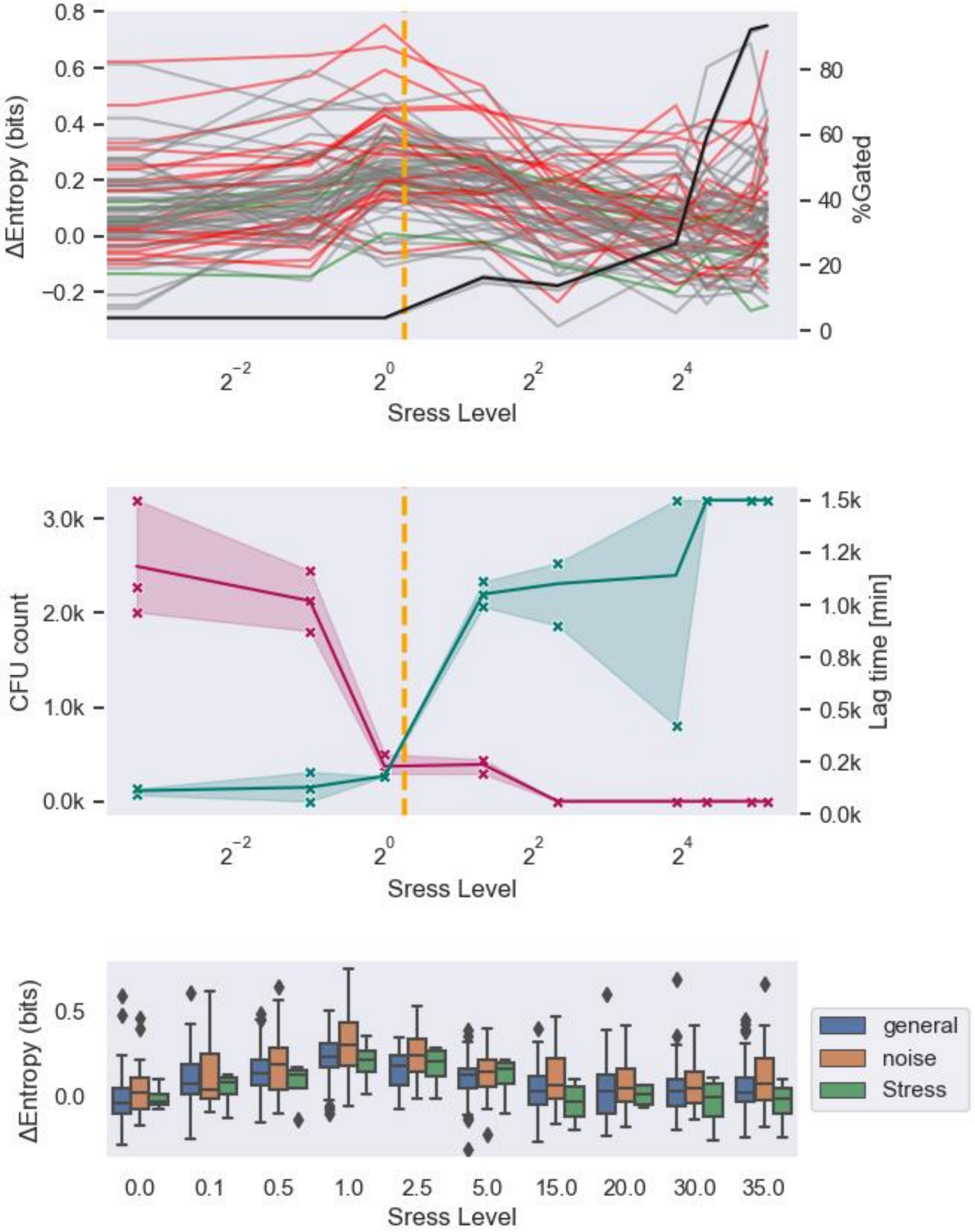

**Figure S7: Entropy of gene expression and stress parameters for different concentrations of $H_2O_2$.** The top panel shows the entropy of gene expression for all genes, where "noisy" genes are shown in red and stress genes in green. The black line shows the % of dead cells as measured using live/dead far-red stain. The middle panel shows stress quantification via CFU and recovery lag time. The dashed orange line signifies the end of the physiological range according to Yeast Stress responses (4). The bottom panel shows the entropy distribution data as the box plots. Notice that all gene groups display a similar trend.

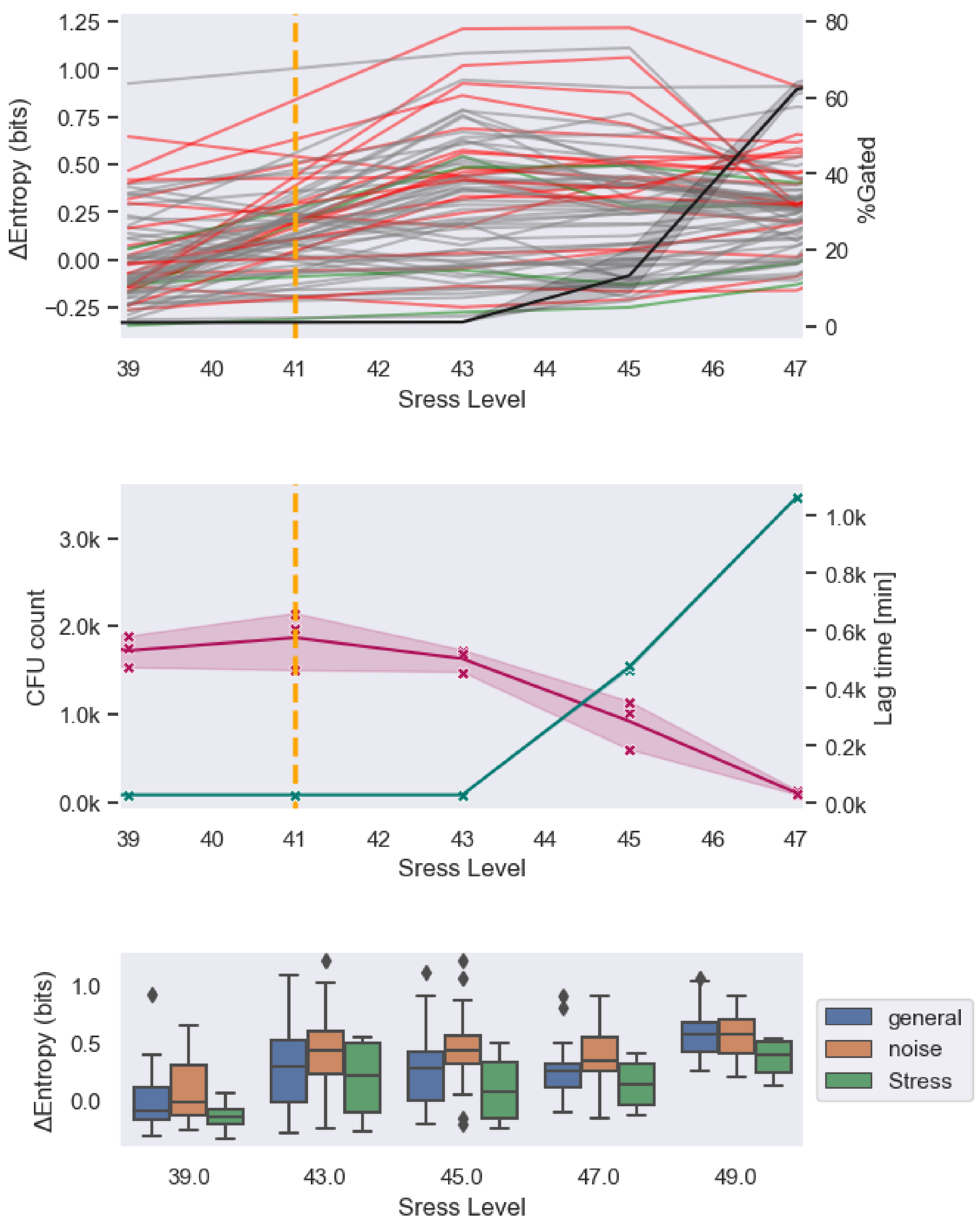


**Figure S8: Entropy of gene expression and stress parameters for different temperatures.** The top panel shows the entropy of gene expression for all genes, where “noisy” genes are shown in red and stress genes in green. The black line shows the % of dead cells as measured using live/dead far-red stain. The middle panel shows stress quantification via CFU and recovery lag time. The dashed orange line signifies the end of the physiological range according to Yeast Stress responses (4). The bottom panel shows the entropy distribution data as the box plots. Notice that all gene groups display a similar trend.

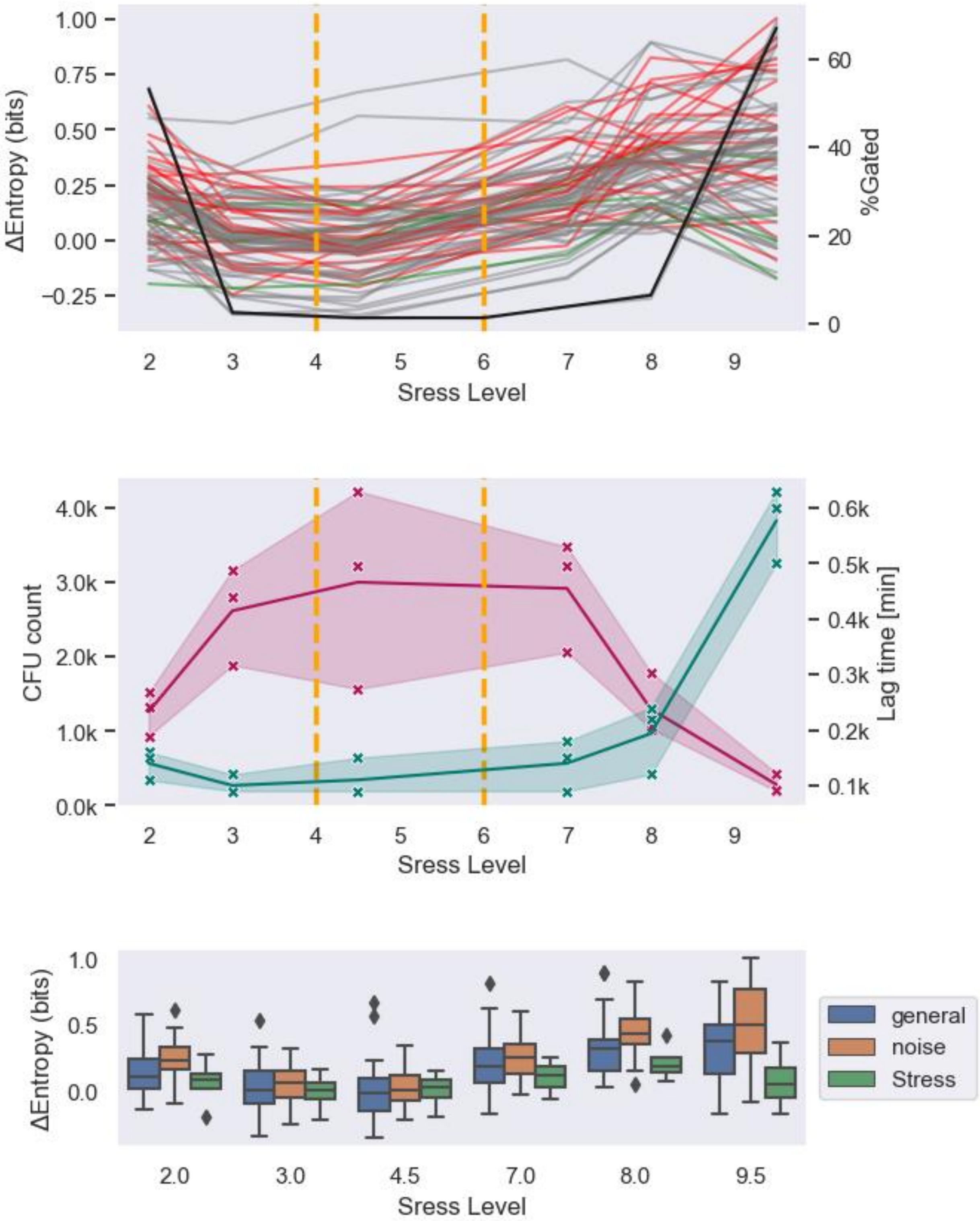

**Figure S9: Entropy of gene expression and stress parameters for different pH levels.** The top panel shows the entropy of gene expression for all genes, where "noisy" genes are shown in red and stress genes in green. The black line shows the % of dead cells as measured using live/dead far-red stain. The middle panel shows stress quantification via CFU and recovery lag time. The dashed orange line signifies the end of the physiological range according to Yeast Stress responses(4). The bottom panel shows the entropy distribution data as the box plots. Notice that all gene groups display a similar trend.


1. Bar-Even A, Paulsson J, Maheshri N, Carmi M, O'Shea E, Pilpel Y, et al. Noise in protein expression scales with natural protein abundance. Nat Genet. 2006 Jun;38(6):636–43.

2. Blake WJ, Kærn M, Cantor CR, Collins JJ. Noise in eukaryotic gene expression. Nature. 2003 Apr 10;422(6932):633–7.

3. Keren L, van Dijk D, Weingarten-Gabbay S, Davidi D, Jona G, Weinberger A, et al. Noise in gene expression is coupled to growth rate. Genome Res. 2015 Dec;25(12):1893–902.

4. Hohmann S. Yeast Stress Responses. Springer Science & Business Media; 2003. 398 p.
